# Glia actively sculpt sensory neurons by controlled phagocytosis to tune animal behavior

**DOI:** 10.1101/2020.11.11.378893

**Authors:** Stephan Raiders, Erik Calvin Black, Andrea Bae, Stephen MacFarlane, Shai Shaham, Aakanksha Singhvi

## Abstract

Glia in the central nervous system engulf neuron fragments during synapse remodeling and recycling of photoreceptor outer-segments. Whether glia passively clear shed neuronal debris, or actively remove neuron fragments is unknown. How pruning of single-neuron endings impacts animal behavior is also unclear. Here we report that adult *C. elegans* AMsh glia engulf sensory endings of the AFD thermosensory neuron. Engulfment is regulated by temperature, AFD’s sensory input, and tracks AFD activity. Phosphatidylserine (PS) flippase TAT-1/ATP8A, functions with glial PS-receptor PSR-1/PSR and PAT-2/α-integrin to initiate engulfment. Glial CED-10/Rac1 GTPase, acting through a conserved GEF complex, executes phagocytosis using the actin-remodeler WSP-1/nWASp and the membrane-sealing factor EFF-1 fusogen. CED-10 levels determine engulfment rates, and engulfment-defective mutants exhibit altered AFD-ending shape and thermosensory behavior. Our findings reveal a molecular pathway underpinning glia-dependent phagocytosis in a peripheral sense-organ, and demonstrate that glia actively engulf neuron-fragments, with profound consequences on neuron shape and animal behavior.

## INTRODUCTION

To interpret its environment accurately and respond with appropriate behaviors, an animal’s nervous system needs to faithfully transmit information from the periphery and through neuron-neuron contacts within the neural network (Bargmann and Marder, 2013). Precision in this information transfer and processing depends partly on neuron receptive-endings (NREs), specialized sub-cellular structures where a neuron receives input from either the external environment or other neurons (Bourne and Harris, 2008; Harms and Dunaevsky, 2007; Shaham, 2010; Singhvi et al., 2016). In the peripheral nervous system (PNS), sensory NREs house the sensory transduction machinery, and appropriate NRE shape dictates sensory information capture. In the central nervous system (CNS), the size and number of interneuron NREs (dendritic spines) help determine the connectome and thereby path of information transfer (Bargmann and Marder, 2013; Eroglu and Barres, 2010; Nimchinsky et al., 2002). While remodeling of NRE shape has been suggested important for experiential learning and memory (Bourne and Harris, 2008; Harms and Dunaevsky, 2007), directly correlating these subcellular changes with animal behavior has been challenging.

Glia are the other major cell-type of the nervous system and approximate neurons in number (von Bartheld et al., 2016). They have been proposed to actively modulate development, homeostasis and remodeling of neural circuits, and have been suggested to influence NRE shape and numbers (Allen and Eroglu, 2017; Stogsdill and Eroglu, 2017; Zuchero and Barres, 2015). One mechanism by which glia may do so is by engulfment of neuron fragments and NREs (Freeman, 2015; Schafer and Stevens, 2013; Wilton et al., 2019), and aberrant neuron fragment uptake by glia is implicated in neuro-developmental as well as neuro-degenerative diseases including Alzheimer’s dementia, Autism and Epilepsy (Chung et al., 2015; Henstridge et al., 2019; Neniskyte and Gross, 2017; Schafer and Stevens, 2013; Vilalta and Brown, 2018; Wilton et al., 2019).

Nonetheless, fundamental questions about the roles and mechanisms of glia-dependent phagocytosis remain. One, whether glia initiate engulfment or passively respond to neuron shedding is unclear. Two, correlating glia-dependent remodeling at single synapse or NRE with changes in animal behavior remains impossible in most systems (Koeppen et al., 2018; Wang et al., 2020). Also, glial engulfment mechanisms have been primarily dissected in the context of injury or development, and their impact on adult neural functions remains less understood. Finally, whether glia-dependent engulfment occurs in the peripheral nervous system or dictates normal sensory functions has not yet been shown.

The nervous system of *C. elegans* is comprised of 302 neurons and 56 glia (Singhvi and Shaham, 2019; Sulston et al., 1983; White et al., 1986). These arise from invariant developmental lineages, form invariant glia-neuron contacts, and each neuron performs defined functions to enable specific animal behaviors. These features allow single-cell and molecular analyses of individual glia-neuron interactions with exquisite precision(Singhvi et al., 2016; Singhvi and Shaham, 2019). Here, we describe our discovery that the *C. elegans* AMsh glial cell engulfs NRE fragments of the major thermosensory neuron of the animal, AFD. Thus, this critical glial function is conserved in the nematode and across sense-organ glia. Engulfment rates are regulated by temperature and track AFD neuron activity. Engulfment requires the phospholipid transporter TAT-1/ATP8A, α-integrin PAT-2, and glial phosphatidylserine receptor PSR-1. PSR-1 engages a conserved ternary GEF complex (CED-2/CrkII, CED-5/DOCK180, CED-12/ELMO1) to activate CED-10/Rac1 GTPase. Downstream effects include the actin remodeling factor WSP-1/nWASp and EFF-1/fusogen which mediates phagocytic cup sealing. Importantly, glial CED-10/Rac1 expression levels dictate NRE engulfment rates, and perturbation of glial engulfment leads to defects in AFD-NRE shape and associated thermosensory behavior.

Taken together, our studies show that glia actively regulate engulfment rather than passively clearing shed neuron debris and do so by repurposing components of the apoptotic phagocytosis machinery. However, whereas cell corpse engulfment is an all-or-none process, glia-dependent engulfment of AFD endings is dynamic. Our findings directly link pruning of a single glia-neuron interaction site to animal behavior. We propose that other glia may similarly deploy regulated phagocytosis to tune sensory NREs and synapses, and to dynamically modulate adult animal behaviors.

## RESULTS

### *C. elegans* glia engulf fragments of the AFD neuron receptive-ending

*C. elegans* is an outstanding setting in which to explore how activities of single neurons affect animal behavior and has the potential to shed light on how single-cell pruning events contribute to behavioral plasticity. Sensory-organ and brain-associated glia of the nematode *C. elegans* share molecular, morphological, anatomical, and functional features with vertebrate sense-organ glia and astrocytes, respectively(Bacaj et al., 2008; Katz et al., 2018; Katz et al., 2019; Singhvi and Shaham, 2019; Wallace et al., 2016). To uncover molecular mechanisms driving glia-dependent NRE uptake, we investigated interactions between the *C. elegans* AMsh glial cell and the NRE of the temperature-sensing neuron AFD (Singhvi et al., 2016; Singhvi and Shaham, 2019; Wallace et al., 2016). The AFD NRE is comprised of ~45 microvilli and a single cilium and is embedded in the AMsh glial cell. An adherens junction between the AFD NRE base and the AMsh glial cell serves to isolate this compartment (Figure 1A,B) (Doroquez et al., 2014; Perkins et al., 1986).

**Figure 1.**
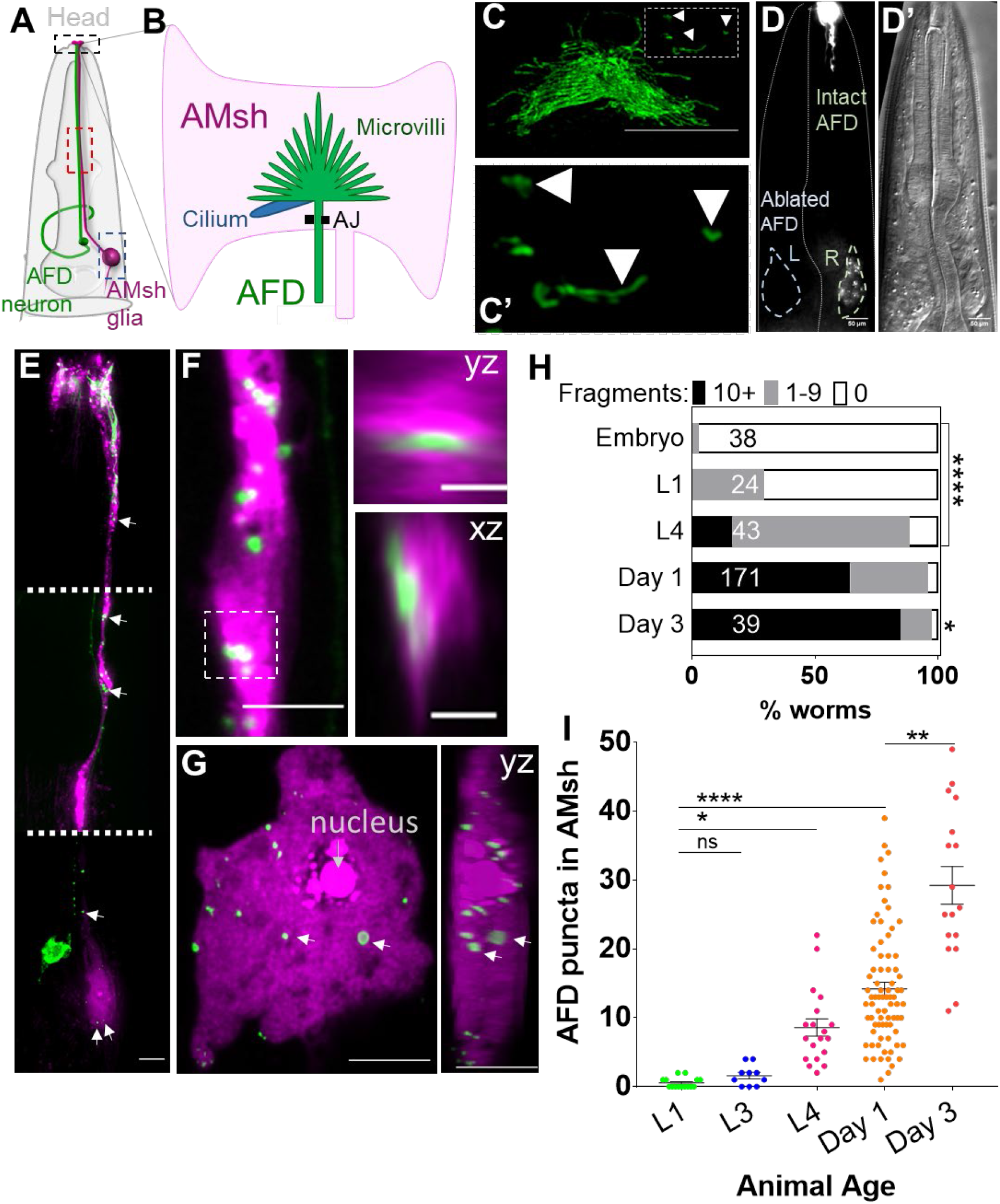
AMsh glia engulf fragments of AFD-NRE. **(A)** Schematic of a *C. elegans* head depicting AFD neuron and AMsh glial cell body and processes. Anterior is to the top. Black box: zoomed in Figure 1B,C; red box zoomed in Figure 1F; blue box zoomed in Figure 1G. **(B)** The AMsh glia’s anterior ending ensheathes AFD-NRE dendrite, which comprises ~45 microvilli (green) and a single cilium (blue). AJ= adherens junction between AMsh glia and AFD neuron **(C, C’)** P_SRTX-1b_:SRTX-1:GFP specifically labels AFD-NRE microvilli. Arrows indicate microvilli fragments disconnected from the main AFD-NRE structure, zoomed in C’. Anterior to top. Scale bar: 5μm. **(D, D’)** Day 1 adult animal with left AFD neuron ablated by laser-microsurgery during L1 larval stage. Left AMsh soma (blue outline) lacks AFD-NRE fragments, right AMsh soma (green outline) contains fragments. D, fluorescence micrograph; D’, DIC image **(E-G)** AMsh glia (magenta) show AFD-NRE puncta throughout the cell (E) including the process (F) and soma (G). Image in (E) is a composite of three exposure settings of a single animal, stitched where indicated by dotted white line. Orthogonal slices of AMsh glial process (F, Scale bar: 2μm) and cell body (G) show AFD-NRE fragments completely within AMsh glia. Scale bar: 5μm. **(H)** Population scores of wild type animals with AFD-NRE labeled fragments within AMsh soma at different developmental stages. Puncta numbers are quantified into three bins (≥10 fragments, black bar), (1-9 fragments, grey bar), (0 fragments, white bar). See Methods for details. N= number of animals. X-axis: percent animals with fragments. Y-axis: developmental stage **(I)** Quantification of AFD-NRE labeled fragments within AMsh soma at different developmental stages. X-axis: Developmental stage, Y-axis: number of puncta per AMsh glial cell-soma. Median puncta counts and N (number of animals): L1 Larva (0.5±0.2 puncta, n=15 animals), L3 Larva (1.6±0.5 puncta, n=10 animals), L4 Larva (8.5±0.2 puncta, n=19 animals), Day 1 Adult (14±1 puncta, n=54 animals), Day 3 Adult (29±3 puncta, n=17 animals) Statistics: Kruskal-Wallis H test w/ multiple comparison. P<0.05 =*, P<0.005 = **, P<0.0005 = ***, P<0.00005 = **** ns = p>0.05.

Upon imaging AFD NREs in transgenic animals labeled with any of five different fluorescent reporter proteins, we consistently observed labeled fragments disconnected from the neuron (Figure 1C; Figure S1A, S1B, Movie 1). Surprisingly, two-color imaging revealed that many of these fragments reside within the AMsh glial cell body and process (Figure 1E-G). Population distribution analyses revealed that ~65% of animals have AMsh glia containing >10 puncta and another ~32% animals have 1-9 puncta/glia (n=171) (Figure 1H) (see Methods for binning details). Each single AMsh glial cell of 1-day-old adult animals raised at 20°C has an average of 14 ± 1 puncta (n=54) (Figure 1I). Analyses of 3D super-resolution microscopy images reveal that the size of individual glial puncta average 541 ± 145 nm along their long (yz) axis (Figure S1C). These fragments are considerably smaller than recently described exophers extruded from neurons exposed to cellular stress (~3.8 μm in diameter) (Chung et al., 2013; Melentijevic et al., 2017).

### AMsh glia engulf AFD-NRE microvilli

AFD fragments could arise from either the AFD-NRE cilium or microvilli. To distinguish which organelle was engulfed, we undertook two approaches. First, we labeled each organelle with specific fluorescent tags and examined which staining was found in AMsh glia. To probe microvilli, we examined transgenic animals with tagged AFD-microvilli specific proteins SRTX-1, GCY-8, GCY-18 and GCY-23 (Colosimo et al., 2004; Inada et al., 2006). We found that all four transgenic strains consistently show fluorescent puncta in glia (Figure 1C-C’, S1A). To label cilia, we generated transgenic animals with fluorescently tagged DYF-11/TRAF31B1 expressed under an AFD-specific promoter. DYF-11 expresses in all ciliated neurons including the amphids and is an early component of cilia assembly (Bacaj et al., 2008; Kunitomo and Iino, 2008; Perkins et al., 1986; Starich et al., 1995). DYF-11:GFP localizes to AFD cilia (Figure S2A) but we found no DYF-11:GFP puncta in AMsh glia (Figure S1A). Second, we examined mutants lacking microvilli or cilia. The development of AFD microvilli requires the terminal selector transcription factor TTX-1/Otx1/Orthodenticle (Hobert, 2016; Satterlee et al., 2001). AFD NRE puncta are absent from AMsh glia of *ttx-1(p767)* mutants which lack microvilli but are still present in *dyf-11(mn392)* mutants (Figure 2A-C). These data indicate that the observed puncta in AMsh glia derive from AFD microvilli.

**Figure 2.**
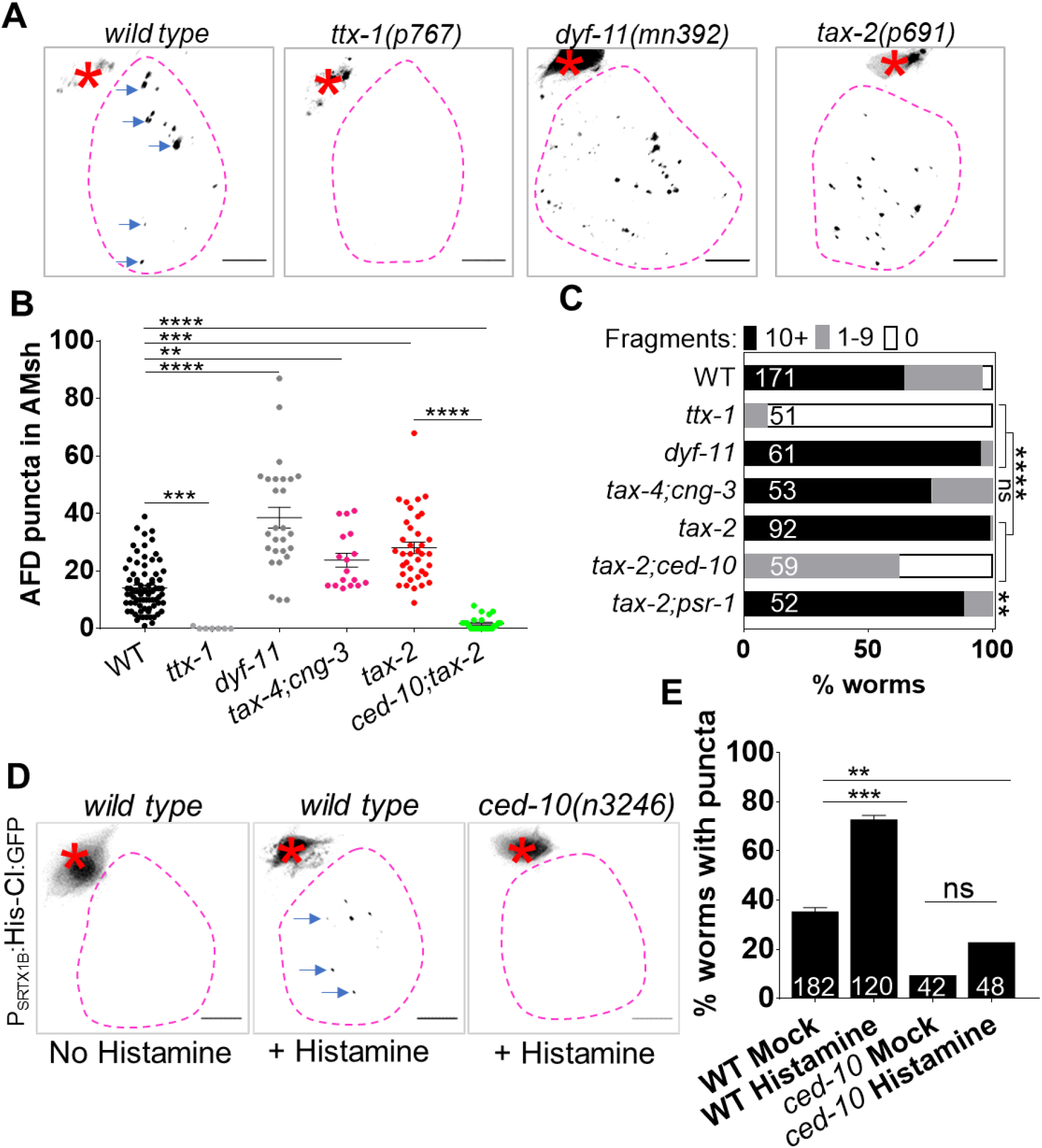
AMsh glia engulf AFD-NRE microvilli in activity-dependent manner. **(A)** Panel showing AFD-NRE tagged puncta (blue arrows) within AMsh glial cell soma (magenta outline) in different genetic backgrounds as noted. AFD cell-body (red Asterix). Scale bar: 5μm. **(B)** Quantification of puncta within AMsh cell soma in microvilli, cilia or activity mutants. Refer Figure 1I for data presentation details. Median puncta counts and N (number of animals): *wild type* (14±1 puncta, n=54 animals), *ttx-1(p767)* (0±1 puncta, n=7 animals), *dyf-11(mn392)* (40±2 puncta, n=27 animals), *tax-2(p691)* (28±2 puncta, n=37 animals), *tax-4(p678);cng-3(jh113)* double mutants (24±2 puncta, n=17 animals), *ced-10(n3246);tax-2(p691)* mutants (2±0.5 puncta, n=25 animals) **(c)** Population counts of animals with AMsh glial puncta. Refer Figure 1H for data presentation details. Alleles used: *ttx-1(p767)*, *dyf-11(mn392)*, *tax-4(p678);cng-3(jh113)*, *tax-2(p691)*, *ced-10(n3246);tax-2(p691)*, *tax-2(p691);psr-1(tm469)*. **(D)** Panel depicts AFD-labeled puncta (blue arrows) within AMsh glial cell soma (magenta outline) in different experimental conditions/genotype as noted. Scale bar: 5μm. **(E)** Percent *wild type* or *ced-10(n3246)* mutant animals with observable GFP+ puncta with or without histamine. N= number of animals.

Time-lapse microscopy supports the notion that the fragments we observed originate from the AFD NRE microvilli (Figure S1A, B; Movie 1). In 1-day-old adults at 20°C, individual puncta, as labeled by microvilli specific markers, separate from the NRE at a frequency of 0.8 ± 0.3 events/minute, and travel at 1.05 ± 0.1 μm/sec down the glial process towards the cell body, consistent with motor-protein-dependent retrograde trafficking (Figure S1B; Movie 1) (Maday et al., 2014; Paschal et al., 1987). The cylinder diameter of AFD microvilli, measured by electron microscopy (EM), is 214 ± 30 nm (Figure S1D,E), suggesting that puncta may be microvilli fragments of varying lengths.

Upon ablation of one of the two bilateral AFD neurons by laser microsurgery in first larval-stage (L1) animals, we find that fragment formation is blocked on the operated but not on the un-operated side, or in mock-ablated animals (Figure 1D; Figure S1F). Similar results are seen with stochastic genetic ablation of AFD with the pro-apoptotic BH3-domain protein EGL-1, expressed using an AFD-specific embryonic-onset promoter (Figure S1G). Based on these findings, we conclude, that AMsh glia engulf fragments of the AFD NRE microvilli in *C. elegans*.

### AMsh glial engulfment of AFD-NRE occurs in adult animals post-development

Engulfment of neuronal fragments by glia has been suggested to refine neuronal circuit connectivity in development, following completion of nervous system xassembly(Wilton et al., 2019). To determine when *C. elegans* AMsh glia initiate engulfment of AFD NRE fragments, we counted glia-engulfed NRE puncta at different life stages. We found that these puncta are rarely found in embryos or early larval stages, but are easily detected in L4 larvae, and increase in numbers during adulthood (Figure 1H, I). Thus, consistent with our L1 laser ablation studies above, engulfment of AFD NREs by glia occurs continually post-development.

### Glial engulfment tracks neuron activity

Detailed quantification of puncta accumulation in AMsh glia of *dyf-11* mutants (that lack AFD-NRE cilia) revealed, in fact, an increased puncta number within AMsh glia compared to wild type animals (38 ± 3, n=27, vs. 14 ± 1, n=54, in 1-day-old mutant and wild-type adults, respectively, at 20°C) (Figure 2B). A larger fraction of *dyf-11* mutant animals exhibiting >10 puncta/glia (95%) compared to wild type animals (64%) (Figure 2C). Thus, glia engulf more fragments in animals with defective cilia. Previous studies had shown that cyclic-nucleotide-gated (CNG) ion channels localize to the AFD cilium base and are required for AFD neuron firing in response to temperature stimuli. Importantly, these channels are mis-localized in cilia-defective mutants(Nguyen et al., 2014; Satterlee et al., 2004), and cilia-defective mutants exhibit deficits in thermotaxis behavior(Tan et al., 2007). We therefore wondered whether supernumerary puncta in *dyf-11* mutants arise from defects in neuron activity.

To test this, we counted puncta in animals deficient for either TAX-2, the sole β-subunit, or TAX-4 and CNG-3, α-subunits of AFD-expressed CNG channels(Cho et al., 2004; Hellman and Shen, 2011; Satterlee et al., 2004). We found that both *tax-2* single mutants (28 ± 2 puncta, n=37) and *tax-4; cng-3* double mutants (24 ± 2.3 puncta, n=17) accumulate more puncta per glial cell than wild-type animals (Figure 2A,B), and a larger fraction of the animal population had 10+ puncta (*tax-2,* 99%*; wt,* 64%) (Figure 2C). Furthermore, we noted that the number of puncta in glia tracks cultivation temperature for the animal (Figure S2B), possibly reflecting on the level of neuron stimulation.

To test whether activity-dependent regulation of glial puncta accumulation is dynamic, we engineered transgenic animals expressing a histamine-gated chloride channel in an otherwise wild-type background(Pokala et al., 2014). Acute silencing of AFD by histamine addition to the cultivation media led to puncta enrichment in AMsh glia within 24 hours in Day 1 adult animals (Figure 2D-E). Taken together, these results suggest that AMsh glia dynamically repress NRE engulfment based on AFD neuron activity post-neural development.

### The PS receptor PSR-1 mediates glial engulfment

To uncover the molecular machinery regulating AFD NRE engulfment by AMsh glia, we examined mutants in components required for apoptotic cell engulfment in *C. elegans* (Figure S3A). Surprisingly, loss of the conserved engulfment receptor CED-1/Draper/ MEGF10(Mangahas and Zhou, 2005), which is also required for glia-mediated removal of neuron debris in other contexts(Cherra and Jin, 2016; Nichols et al., 2016), does not block NRE fragment uptake (Figure 3A). Removal of the CED-1 effectors, CED-6/GULP or CED-7/ABCA1(Flannagan et al., 2012; Reddien and Horvitz, 2004; Zhou et al., 2001), also does not block engulfment (Figure 3A). Like *Drosophila,* the *C. elegans* genome does not encode a direct ortholog of the TAM MeRTK, which recognizes neuronal debris in vertebrates. We tested related VEGF and FGF receptor tyrosine kinases, and these also seem not to be required (Figure S3B) (Popovici, 1999). Thus, some of the major components driving apoptotic cell recognition and phagocytosis are not required for AFD NRE engulfment.

**Figure 3.**
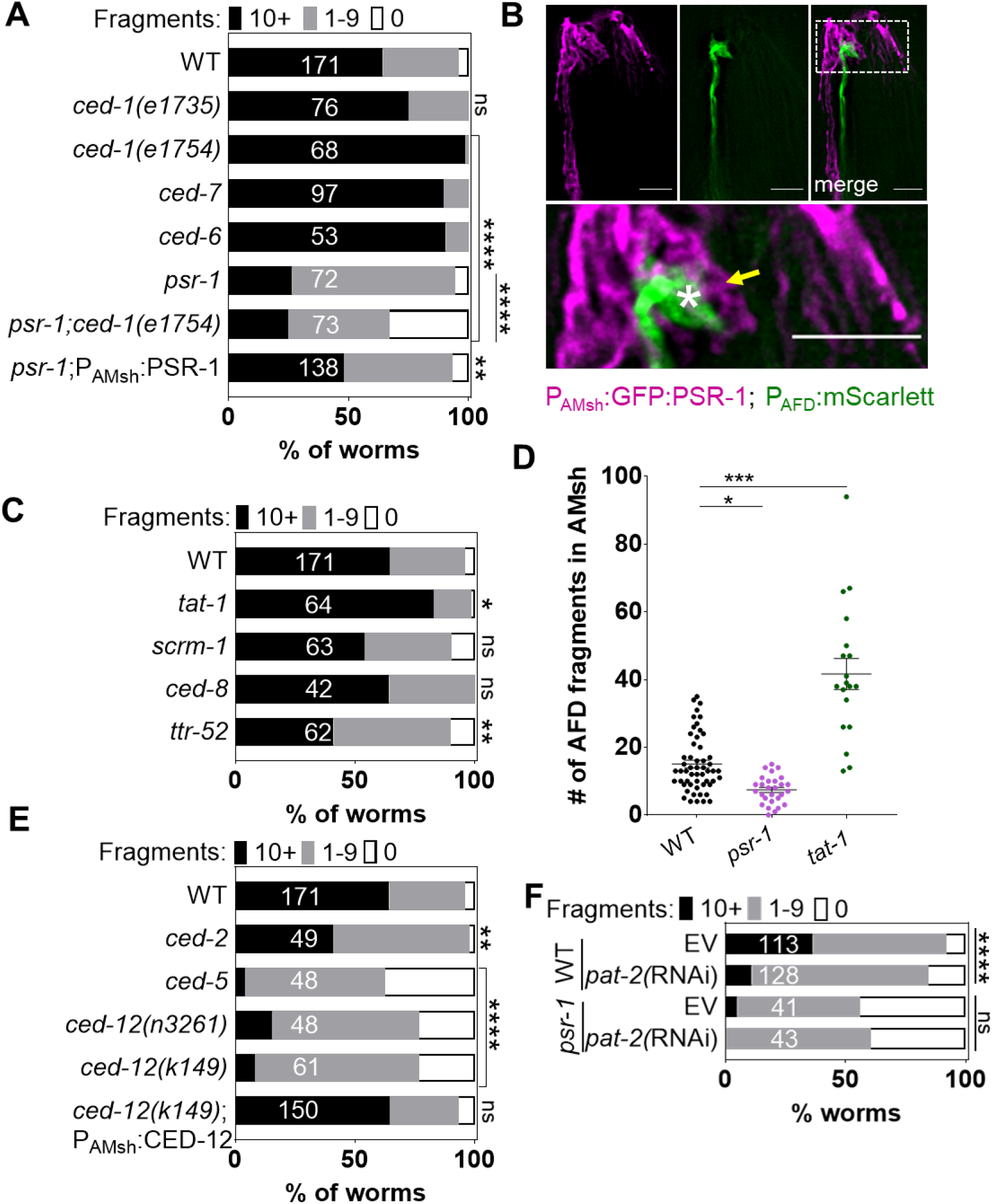
Engulfment of AFD-NRE by AMsh glia requires the phosphatidylserine receptor PSR-1 and integrins. **(A,C,E,F)** Population counts of animals with AMsh glial puncta in genetic backgrounds indicated. Refer Figure 1H for data presentation details. **(A)** Alleles used: *ced-1(e1735),ced-1(e1754),ced-7(n2094),ced-6(n1813),psr-1(tm469)*. **(B)** Fluorescence micrograph of a transgenic animal with GFP tagged PSR-1 expressed specifically in AMsh glia (magenta) localizing on the apical membrane around AFD-NRE (green). GFP:PSR localizes to apical membrane in AMsh glia (yellow arrow) around AFD-NRE (asterisk). Scale bar: 5μm. Bottom: Zoom of box in two-color merged image. **(C)** Alleles used: *tat-1(tm1034), scrm-1(tm805),ced-8(n1819), ttr-52(tm2078)*. Refer Figure 1H for data presentation details. **(D)** Quantification of puncta within AMsh cell soma in phagocytosis pathway mutants. Refer Figure 1I for data presentation details. Median puncta counts and N (number of animals): *wild type* (14±1 puncta, n=54 animals), *psr-1(tm469)* (7.4±0.7 puncta, n=28 animals), *tat-1* (42±5 puncta, n=19 animals)**. (E)** Alleles used: *ced-2(1752)*, *ced-5(n1812)*, *ced12(n3261),* ced*-12(k149)*. Refer Figure 1H for data presentation details. **(F)** RNAi (control *pat-2)* in wild-type or *psr-1(tm469)* mutant animals.

Loss of *psr-1*, which encodes a phosphatidylserine receptor, was previously reported to have modest effects on apoptotic cell corpse engulfment in *C. elegans* and zebrafish (Hong et al., 2004; Wang et al., 2003). In contrast, we found that deletion of *psr-1* dramatically reduces AFD NRE uptake by AMsh glia (Figure 3A, D). Expression of the PSR-1C isoform in AMsh glia significantly rescues *psr-1* mutant defects (Figure 3A), suggesting that PSR-1 functions within glia for uptake. To examine PSR-1 localization, we generated transgenic animals carrying tagged GFP:PSR-1 expressed in AMsh glia, and found that the fusion protein localizes to membranes, including those around AFD NRE microvilli (Figure 3B). Together, these results strongly suggest that NRE fragments are recognized by glial PSR-1 for engulfment.

Loss of *psr-1* significantly, but only partially, curbs excess engulfment observed in animals lacking the TAX-2 CNG channel subunit (Figure 2C). This observation suggests two things. One, that PSR-1 likely mediates engulfment downstream of neuron activity. Second, that additional engulfment receptors may be at play. Consistent with this, although *psr-1* deletion reduces NRE engulfment by AMsh glia significantly, uptake of neuronal fragments is not completely eliminated (Figure 3A). We therefore wondered if CED-1/MEGF10/Draper1 functions redundantly with PSR-1. However, engulfment defects of *psr-1* mutant animals are not enhanced by a mutation in *ced-1* (Figure 3A). This suggests that *psr-1* and *ced-1* are unlikely to function redundantly and corroborates that *ced-1* is dispensable for this process.

### Integrins and the PS-bridging molecule TTR-52 regulate glial engulfment

We wondered which other receptor might function with PSR-1 to mediate engulfment. The INA-1 α-integrin subunit is implicated in apoptotic cell phagocytosis in *C. elegans*, and animals carrying the *ina-1(gm144)* mutation accumulate cell corpses(Hsieh et al., 2012; Neukomm et al., 2014; Saenz-Narciso et al., 2016). Integrins have also been implicated in engulfment by retinal RPE glia (Mao and Finnemann, 2012). We therefore tested a role for integrins in our setting and found that while this mutation has no effect on NRE engulfment (Figure S3B), loss of the only other *C. elegans* α-integrin subunit, PAT-2, triggered by RNA interference (RNAi), significantly blocks AMsh glial engulfment of the AFD NRE in wild-type animals (Figure 3F). Furthermore, PAT-2 loss suppresses enhanced engulfment in animals lacking *tax-2*, as well as in *tax-2; psr-1* double mutants (Figure S2C). Finally, the α-integrin INA-1 is activated for cell corpse engulfment by the opsonin transthyretin protein TTR-11, which binds phosphatidyl serine(Hisamoto et al., 2018). We found that a mutation in *ttr-52*, encoding a different transthyretin protein, reduces NRE uptake (Figure 3C). Thus, PAT-2 integrin, triggered perhaps by TTR-52, functions in parallel to PSR-1 to promote NRE engulfment.

### The phospholipid transporter TAT-1 regulates glial engulfment

The involvement of PSR-1, PAT-2/α-Integrin, and TTR-52 in AFD NRE uptake by AMsh glia suggests that PS exposure on neuron fragments may facilitate their engulfment. The Class IV P-type ATPase ATP8A is a phospholipid transporter which maintains PS in the plasma membrane inner leaflet. A mutation in *tat-1*,(ATP8A ortholog), results in increased PSR-1 dependent apoptotic cell corpse engulfment in *C. elegans*, presumably by increasing PS levels on the outer membrane leaflet(Darland-Ransom et al., 2008)(Darland-Ransom et al., 2008). We found that *tat-1* mutants have increased numbers of engulfed puncta in glia (Figure 3C,D), consistent with PS-presentation underlying NRE engulfment. Mutations affecting the functions of the Xkr8 factor CED-8 or the scramblase SCRM-1/PLSCR, which promote PS-presentation for cell-corpse phagocytosis(Bevers and Williamson, 2016; Wang et al., 2007) do not affect NRE uptake (Figure 3C), suggesting that other mechanisms normally drive PS exposure on AFD NRE fragments.

### Glial Rac1 GTPase CED-10 controls rate of engulfment

How might PSR-1 and PAT-2 integrin promote NRE engulfment? Rac1 GTPase is a known downstream effector of a number of apoptotic phagocytosis pathways (Flannagan et al., 2012; Reddien and Horvitz, 2004; Wang and Yang, 2016). It is also implicated in engulfment of photo-receptor outer segments by RPE glia-like cells in mammals (Kevany and Palczewski, 2010; Nichols et al., 2016). We therefore decided to test the role of *ced-10* in AMsh glial engulfment of AFD-NRE.

We found that two independent mutations in *ced-10* nearly fully block engulfment of AFD NRE fragments by AMsh glia (Figure 4A-D). *ced-10* mutations also suppress increased glial puncta accumulation in *tax-2* mutants and in transgenic animals with chemo-genetically silenced AFD neurons (Figure 2B-E). Expressing CED-10 only in AMsh glia completely restores engulfment to *ced-10* mutants (Figure 4A-C). Strikingly, transgenic expression of CED-10 is also sufficient to overcome the partial loss of NRE engulfment in *psr-1* mutants (Figure 4D). Thus, CED-10/Rac1 functions in AMsh glia downstream of PSR-1 to promote engulfment of AFD NREs.

**Figure 4.**
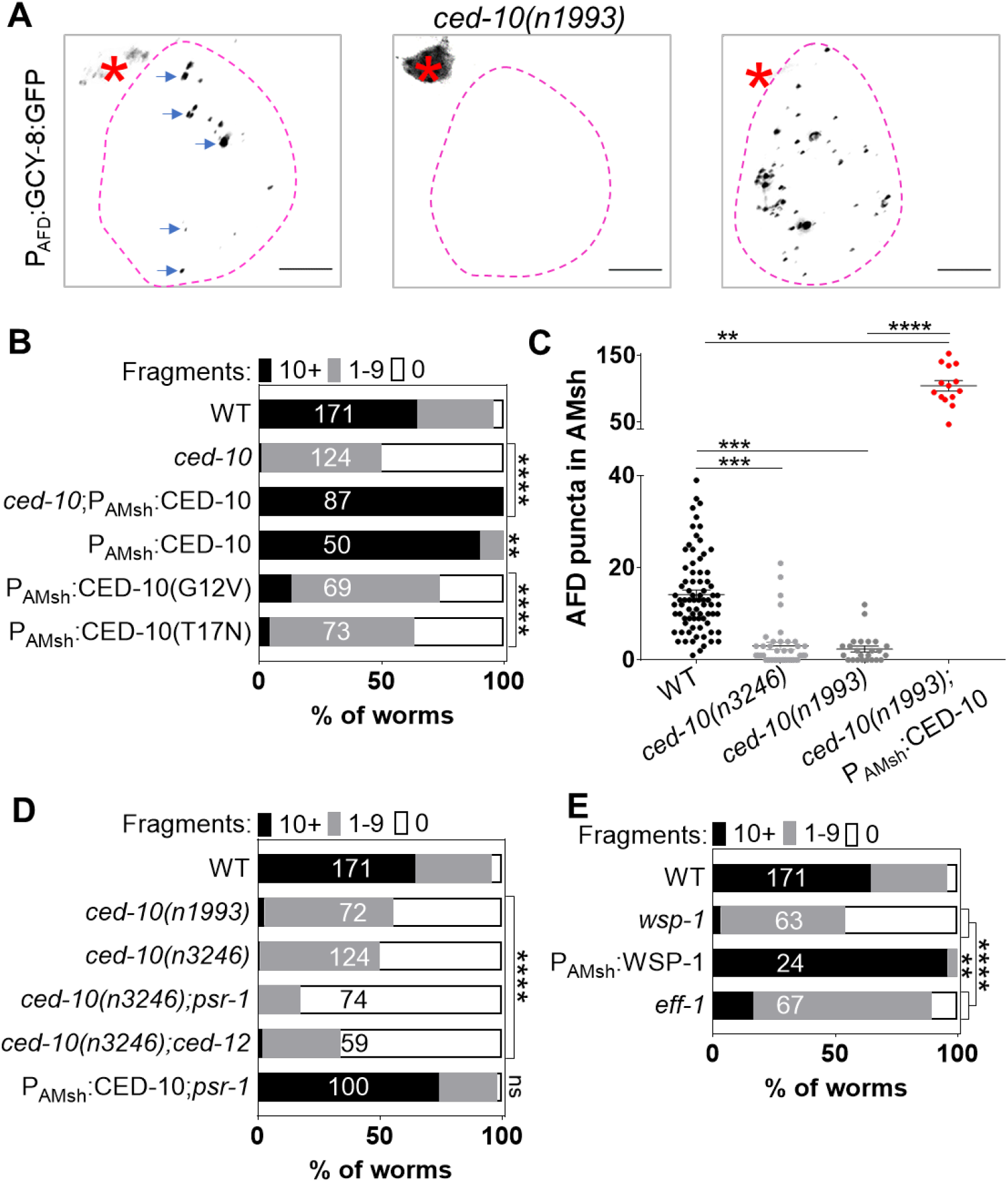
Glial CED-10 levels and actin remodeling control rate of engulfment. **(A)** Panel showing AFD-NRE tagged puncta (blue arrows) within AMsh glial cell soma (magenta outline) in different genetic backgrounds as noted. AFD cell-body (red Asterix). Scale bar: 5μm. **(B,D,E)** Population counts of animals with AMsh glial puncta in genetic backgrounds indicated. Refer Figure 1H for data presentation details. **(B)** Alleles used if not otherwise noted: *ced-10(n1993)*. CED-10(G12V) and CED-10(T17N) is a constitutively active or dominant negative form of CED-10, respectively. **(C)** Quantification of puncta within AMsh cell soma in phagocytosis pathway mutants. Refer Figure 1I for data presentation details. Median puncta counts and N (number of animals): *wild type* (14±1 puncta, n=54 animals), *ced-10(n3246)* (3±1 puncta, n=39 animals), *ced-10(n1993)* (2.3±0.5 puncta, n=24 animals), *ced-10(n1993);* P_AMsh_:CED-10 (105±8 puncta, n=14 animals). **(D)** Alleles used: *ced-10(n1993), ced-10(n3246), psr-1(tm469), ced-12(k149)*. Refer Figure 1H for data presentation details. **(E)** Alleles used: *eff-1(ns634), wsp-1(gm324)*. Refer Figure 1H for data presentation details.

Our observation that CED-10 overexpression restores engulfment to animals lacking PSR-1 led us to hypothesize that CED-10 activity could be the rate limiting step in NRE engulfment. Indeed, in an otherwise wild-type background, overexpression of CED-10, but not PSR-1 or CED-12, leads to increased NRE engulfment (Figure 3A,E; Figure 4A,B). As expected, overexpression of dominant-negative CED-10^T17N^ blocks engulfment (Figure 4B). We therefore infer that glial CED-10 is necessary and sufficient to regulate the rate at which AMsh glial engulf AFD NRE fragments.

These results lead to two additional and important conclusions. One, because these manipulations are specific to AMsh glia, these studies are consistent with the notion that AMsh glia actively engulf AFD NRE fragments, instead of passively internalizing shed debris. Two, these reveal that glial pruning defects are not simply a secondary consequence correlated with neuronal control of AFD-NRE shape. Rather, glial pruning can control AFD-NRE shape causally. Specifically, we note that while both CED-10 over-expression and *ttx-1* mutants have short NRE (Satterlee et al., 2001) (Figure S4B); unlike animals overexpressing CED-10, *ttx-1* mutants have fewer puncta, not more (Figure 2A). Similarly, while both *ced-10* and *tax-2* mutants have longer, disorganized NRE (Satterlee et al., 2004; Singhvi et al., 2016), *tax-2* mutants have more puncta, not fewer. We propose that glial pruning is one mechanism to control NRE shape in response to activity states. It is likely that this cooperates with other independent neuronal mechanisms to do so.

CED-10 executes phagocytic arm extension by mediating actin remodeling(Wang and Yang, 2016). To test whether it has a similar role during NRE engulfment, we examined animals bearing a loss-of-function mutation in the actin polymerization factor WSP-1. We found that these animals exhibit a block in NRE engulfment (Figure 4E). Further, similar to over-expression of CED-10, we found that levels of WSP-1 also lead to increased NRE engulfment (Figure 4E). Together with the results on *ced-10* above, we infer that CED-10 dependent actin remodeling is the rate limiting step for AMsh glia to engulf AFD-NRE.

### Additional components of the apoptotic phagocytosis machinery mediate engulfment by glia

CED-10 is activated by the CED-2/CrkII, CED-5/DOCK1 and CED-12/ELMO1 ternary GEF complex downstream of PSR-1 during apoptotic cell engulfment, and by UNC-73/Trio Rac-GEF during cell-migration(Lundquist et al.). We found that mutations in *ced-2, ced-5*, or *ced-12* impair NRE engulfment, while loss of *unc-73* does not (Figure 3E; Figure S3B-D). Expression of the CED-12B isoform specifically in AMsh glia is sufficient to rescue *ced-12* mutants defects (Figure 3E).

Furthermore, the *C. elegans* cell-cell fusogen EFF-1 has been implicated in phagosome sealing during phagocytosis of cell process debris(Ghose et al., 2018), and *eff-1* mutants also exhibit defective NRE engulfment (Figure 4E). We conclude that the ternary CED-2/CED-5/CED-12 GEF complex activates CED-10 in AMsh glia for AFD NRE engulfment. CED-10 in turn drives actin rearrangements mediated by WSP-1, which promote cellular extensions around NRE fragments, and which eventually seal using the EFF-1 fusogen.

### Glial engulfment regulates AFD neuron receptive-ending shape and animal behavior

Does glial engulfment of AFD-NRE have physiological consequences for neuron shape and associated animal sensory behavior? To test this, we first examined AFD NRE shape by 3D super-resolution imaging of transgenic animals bearing a tagged reporter that specifically only marks AFD-NRE microvilli. We found that *ced-10* loss of function or over-expression of dominant negative CED-10^T17N^, which both reduce engulfment, results in elongated AFD NRE microvilli (Figure 5A,B; Figure S4A,B). Conversely, overexpressing wild-type CED-10 or GTP-locked CED10^G12V^ shortens microvilli with age (Figure 5A,B; Figure S4A,B). Unexpectedly, CED10^G12V^ overexpression results in fewer NRE puncta within glia (Figure 4B). We suspect, based on the morphological impact of this overexpression, that this may occur because engulfment is so efficient, that we can only visualize the few fragments that remain to be degraded. Thus, these findings show that length of AFD-NRE microvilli is continually and dynamically regulated through AMsh glial engulfment of NRE fragments.

**Figure 5.**
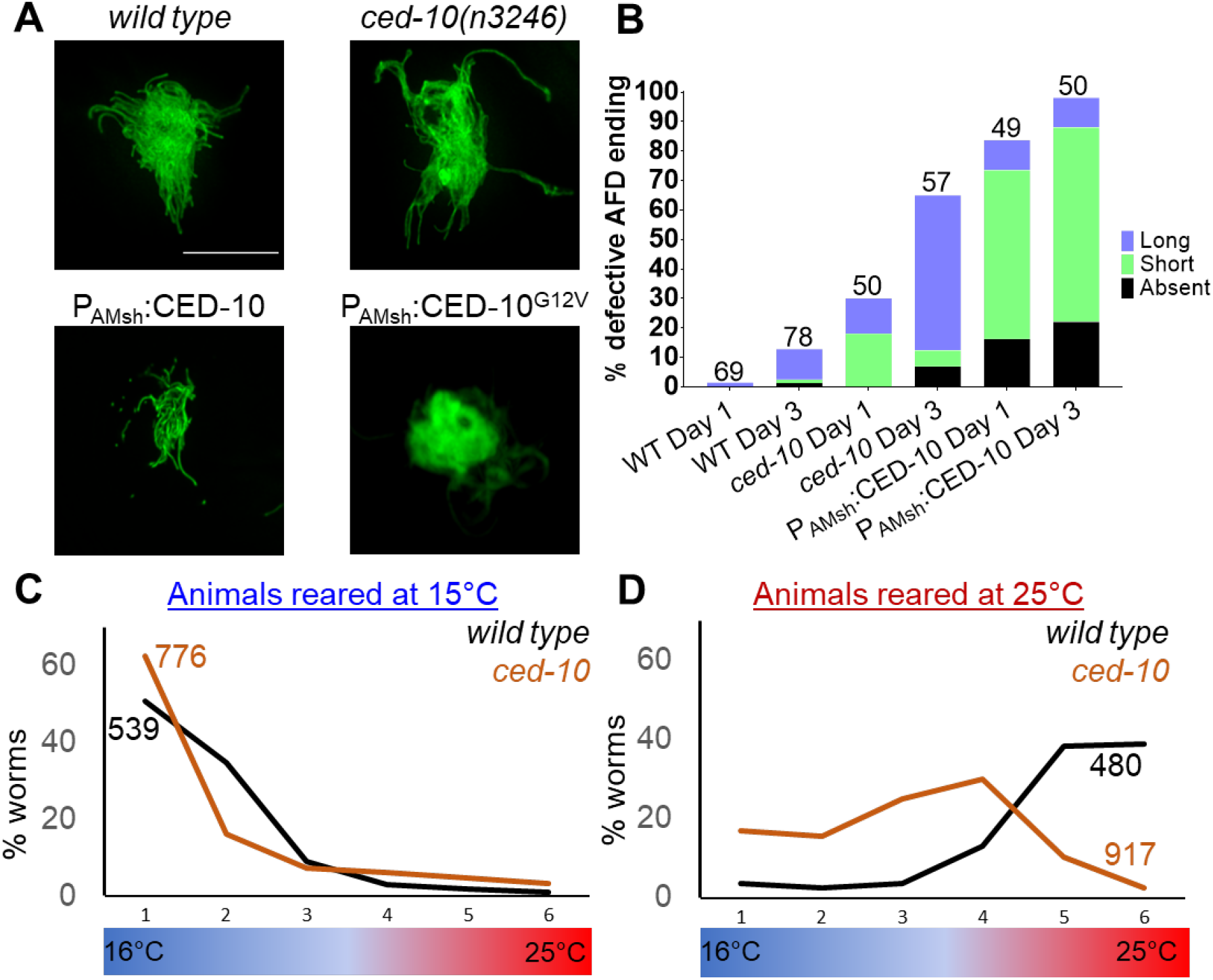
AMsh glial engulfment of AFD-NRE modulates AFD NRE shape. **(A)** AFD-NRE microvilli labelled with GFP in Day 3 adult animals of genotypes as indicated. **(B)** Quantification of percent animals with defective AFD-NRE microvilli shape. N= number of animals scored. **(C,D)** Thermotaxis behavior assays for animals of indicated genotype raised at 15°C **(C)** or 25°C **(D).** Animals assayed 24-hour post-mid-L4 larval stage. N, number of animals.

Does this glia engulfment dependent control of AFD-NRE shape consequently impact associated animal behavior? *C. elegans* migrates to its temperature of cultivation when presented with a linear temperature gradient. This behavior requires receptor guanylyl cyclases on AFD NRE microvilli, which generate cGMP (Hedgecock and Russell, 1975; Mori and Ohshima, 1995; Sengupta and Garrity, 2013). We hypothesized that in *ced-10* mutant animals, the elongated microvilli could house more receptors, resulting in excess thermosensitivity. If so, we reasoned that this should lead animals to migrate to lower temperatures on a thermal gradient. Indeed, we found that while wild type-animals reared at 25°C migrate to their temperature of cultivation, *ced-10* mutants prefer cooler temperatures (Figure 5C,D). This behavior defect is consistent with our previous studies showing that other mutants with increased cGMP levels also exhibit cryophilic behaviors(Singhvi et al., 2016). Thus, glial engulfment of AFD-NRE microvilli, which house thermosensory receptors, regulates its shape and thereby *C. elegans* thermotaxis behavior.

## DISCUSSION

We report our discovery that *C. elegans* glia, like glia of other species, engulf associated neuron endings, highlighting evolutionary conservation of this critical glial function (Figure 6). Exploiting unique features of our experimental model, we demonstrate that glial CED-10 levels dictate engulfment rates, revealing that glia drive neuronal remodeling, and do not just passively clear shed neuronal debris. Indeed, we demonstrate that engulfment is required for dynamic post-developmental maintenance of sensory NRE shape, and behavior. This also extends a role for glial engulfment in the active sensory perception of temperature. Importantly, our studies allow us to directly demonstrate at single-cell resolution that pruning of individual neurons by a single glia modifies animal behavior. This, in conjunction with our finding that phagocytosis is impacted by neuronal activity states, suggests physiological relevance.

**Figure 6.**
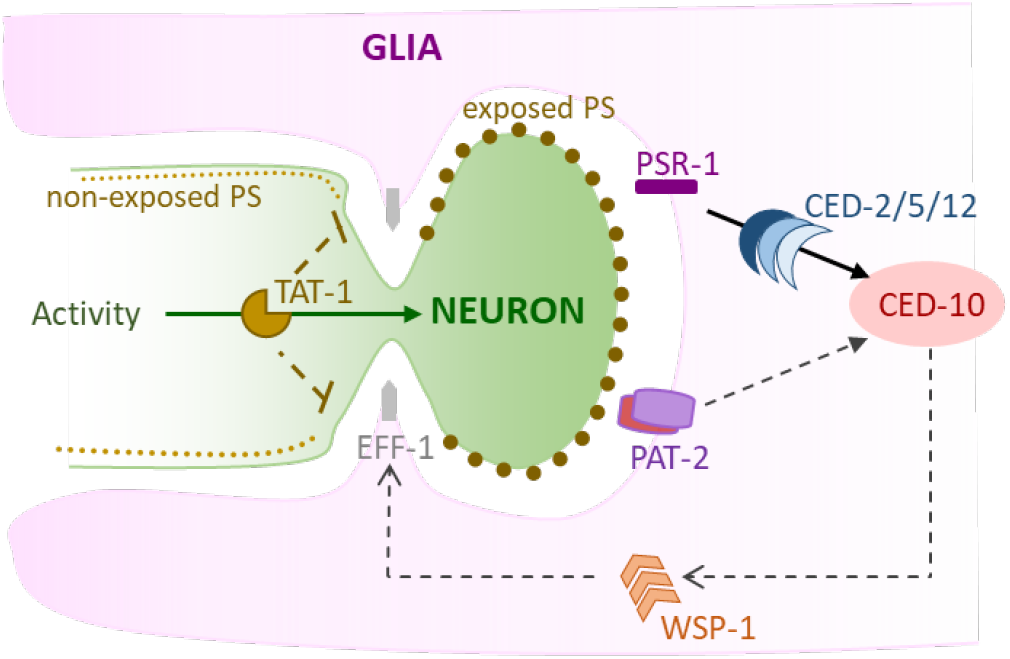
Model of AMsh glial engulfment of AFD NRE. **(A)** Model depicting molecular machinery driving engulfment of AFD neuron microvilli by AMsh glia. TAT-1 maintains phosphatidylserine on the inner plasma leaflet. Neuron activity negatively regulates engulfment. The phosphatidylserine receptor PSR-1 and integrin PAT-2 signal via ternary GEF complex CED-2/5/12 to activate Rac1 GTPase CED-10. CED-10 and its downstream effector, WSP-1, drive engulfment of AFD neuron microvilli fragments. Finally, fusogen protein EFF-1 fuses the forming phagosome to remove microvilli segments.

### Controlled tuning of the phagocytosis machinery

Our studies reveal a fundamental distinction between glia-dependent phagocytosis and other modes of engulfment. Apoptotic cell phagocytosis, glial clearance of injury-induced neuronal debris, and related engulfment events are all or none phenomena: engulfment either occurs or does not. By contrast, we show here that in AMsh glia, engulfment rate is dynamically tuned throughout animal life to modulate NRE morphology, impacting animal behavior. Indeed, to our knowledge this is the first description of phagocytosis machinery being dynamically tuned like a rheostat. The molecular parallels between the engulfment machinery in the peripheral sense-organ AMsh glia, and other CNS glial engulfment leads us to posit that controlled phagocytic machinery may similarly regulate glial engulfment in other settings.

### Distinct receptors mediate PS-dependent glial pruning

Accompanying this more versatile engulfment program is a shift in the relevance of specific engulfment receptors. Apoptotic phagocytosis in *C. elegans* relies predominantly on CED-1, with the PS-receptor PSR-1 playing a minor role (Wang et al., 2003; Wang and Yang, 2016). Surprisingly, while CED-1 is dispensable for pruning by AMsh glia, we identified PSR-1/PS-receptor as a novel regulator of glial pruning. Why do CED-1 and PSR-1 have differing valence in apoptotic phagocytosis and glial pruning? One possibility is that this difference in receptors reflects the size of particles engulfed. Supporting this notion, engulfment of small cell-process debris of the *C. elegans* tail-spike cell is also independent of CED-1 (Ghose et al., 2018).

Why do multiple PS-receptors drive engulfment across glia? Further, while we identified PSR-1 and integrins, other PS-receptors identified in glial pruning include CED-1/MEGF10/Draper, MerTK and GPR56 as well as yet-unidentified receptors (Chung et al., 2013; Freeman, 2015; Hilu-Dadia and Kurant, 2020; Kevany and Palczewski, 2010; Li et al., 2020; Nomura-Komoike et al., 2020; Tasdemir-Yilmaz and Freeman, 2014; Vecino et al., 2016). We speculate that PS-receptor diversity might imply functional heterogeneity in glial engulfment across context.

### Mediators of PS-exposure in *C. elegans* glial pruning

PS exposure has emerged as a classic engulfment signal for both apoptotic phagocytosis and glial pruning, but how this is regulated remains enigmatic. We identify this as a conserved feature in *C. elegans* glial engulfment and implicate the phospholipid transporter TAT-1/ATP8A in this process. TAT-1 is a member of the Type 4 family P4 ATPases, which flip PS from exoplasmic to cytoplasmic membrane leaflets (Andersen et al., 2016). We note that murine P4-ATPases ATP8A1 and ATP8A2 express in the nervous system, and knockout mice exhibit deficient hippocampal learning, sensory deficits, cerebellar ataxia, mental retardation, and spinal cord degeneration, and shortened photoreceptor NRE length (Coleman et al., 2014). Given this intriguing parallel, it will be interesting to probe whether ATP8A similarly modulates glial pruning in mammals.

Further, we identify the PS bridging molecule as a regulator of pruning. It is also implicated in apoptotic phagocytosis and nerve regeneration (Neumann et al., 2015; Wang et al., 2010). Retinal RPE glia and cortical astrocytes also require PS-bridging opsonins (Gas6 and MFGE8) to engulf neuron fragments (Bellesi et al., 2017; Kevany and Palczewski, 2010). Whether all glia require PS-opsonization for pruning remains to be determined.

### Glia direct pruning with sub-cellular precision

Our finding that glia control animal behavior by tuning engulfment suggests that engulfment must proceed with extraordinary specificity, so that behavior is optimal. Indeed, we find that AMsh glia prune AFD NRE with sub-cellular precision. While AFD’s actin-rich microvilli are removed by glia, its adjacent microtubule-based cilium is not. Aberrantly excessive/reduced pruning correlate with disease in mammals, hinting that similar sub-cellular precision in marking fragments/endings for engulfment might be involved(Chung et al., 2015; Wilton et al., 2019). How this precision is regulated will be fascinating to explore.

### Peripheral sense-organ glia prune to regulate sensory behaviors

A role for pruning in normal neural functions has so far been investigated for central nervous system glia, which includes the retinal glia. Peripheral support cell glia of the inner ear are known to activate phagocytosis only in injury settings(Bird et al., 2010). Our studies demonstrate that peripheral sense-organ glia prune associated sensory neuron endings to modulate sensory perception and behaviors. Thus, glial pruning is conserved in both the CNS and PNS and is executed by analogous molecular mechanisms.

### Active pruning versus passive clearance of debris

An outstanding question in understanding the role of glia is whether glia actively prune NREs and neuron fragments, or passively clear shed debris. Our finding that glial CED-10 levels dictate engulfment rates demonstrates that this is actively regulated by glia. That engulfment tracks neuron activity further suggests a physiological role for this process.

Together, these findings reveal glial engulfment as an active regulator of neural functions. Importantly, they directly and causally links pruning of individual neuron endings to animal behavior at single-molecule and -cell resolution. This raises the possibility that engulfment may be a general emergent mechanism by which glia dynamically modulate sensory perception and neural functions, across modalities, systems and species.

## ACKNOWLEDGEMENTS

We thank members of the Singhvi laboratory, Harmit Malik, Jihong Bai and Linda Buck for discussions and comments on the manuscript.

## Funding

This research was conducted while AS was a Glenn Foundation for Medical Research and AFAR Junior Faculty Grant Awardee. This work was funded by a Simons Foundation/SFARI grant (488574) and the Anderson Foundation and the Marco J Heidner Foundation Pilot Fund to AS, and a National Institutes of Health grant (R35NS105094) to SS. Some strains were provided by the CGC, which is funded by NIH Office of Research Infrastructure Programs (P40 OD010440) and some from the National BioResource Project, Japan. Some of the studies were performed at the Fred Hutch Shared Resources Core Facilities.

## Author Contributions

SAR and AS designed, performed and analyzed all experiments and co-wrote the manuscript. SS contributed to initial experimental designs and co-edited the manuscript. EB assisted SAR and AB assisted AS in in strain building. SF performed EM imaging. AS initiated the early phase of this study as a postdoctoral fellow of the American Cancer Society, and The Murray Foundation, in lab of SS.

## Competing Interests

The authors declare no competing interests.

## Data and materials availability

The data and reagents that support the findings of this study are available from the corresponding author upon reasonable request. We apologize to those whose work was not cited due to our oversight or to space considerations.

## Notes

### Competing Interest Statement

The authors have declared no competing interest.

