## Supplemental Materials for "Glia actively sculpt sensory neurons by controlled phagocytosis to tune animal behavior"

**This PDF file includes:**

Materials and Methods

Figures. S1 to S4

Captions for Movie S1

Supplementary Materials Reference List

**Other Supplementary Materials for this manuscript include the following:**

Movie S1

#### MATERIALS AND METHODS

##### Worm methods

*C. elegans* animals were cultured as previously described (Brenner, 1974; Stiernagle, 2006). Bristol N2 strain was used as wild type. For all experiments, animals were raised at $20^{o}$C for at least two generations without starvation, picked as L4 larvae onto fresh plate and assayed 1 day later, unless otherwise noted. Germ-line transformations by micro-injection to generate unstable extra-chromosomal array transgenes were carried out using standard protocols (Mello and Fire, 1995). Integration of extra-chromosomal arrays was performed using UV+ tri-methyl psoralen.

##### Strains and plasmids

Some strains listed below in Sections A and B were sourced from (a) the CGC, funded by NIH Office of Research Infrastructure Programs (P40 OD010440), (b) the International C. elegans Gene Knockout Consortium (C. elegans Gene Knockout Facility at the Oklahoma Medical Research Foundation, funded by the National Institutes of Health; and the C. elegans Reverse Genetics Core Facility at the University of British Columbia, funded by the Canadian Institute for Health Research, Genome Canada, Genome BC, the Michael Smith Foundation, and the National Institutes of Health) and (c) National BioResource Project (NBRP), Japan.

###### A. Mutants

LG1: *tax-2 (p691), ced-12(n3261), ced-12(k149), psr-1(tm469), ced-1(e1754), ced-1(e1735), scrm -1(tm805), aex-5(sa23), kpc-1(gk8), unc-73(e936)*

LG2: *eff-1(ns634)*

LG3: *ttr-52(tm2078), ced-6(n1813), tat-1(tm1034), tax-4(p678), ced-7(n2094), ver-1(ok1738), ver-2(ok897), ina-1(gm144)*

LG4: *ced-10(n3246), ced-10(n1993), ced-2(e1752), ced-5(n1812), unc-31(e928), cng-3(jh113)*

LG5: *ttx-1(p767)*

LGX: *dyf-11(mn392), ced-8(n1819), ver-3(ok891), ver-4(ok1079), egl-15(n484)*

**B. Integrated transgenes**

| Strain ID | Chromosome | Genotype | Reference |
|---|---|---|---|
| *nsIs228* | I | $P_{srtx\text{-}1}$*:GFP* | (Colosimo et al., 2004; Singhvi et al., 2016) |
| *nsIs481, nsIs482, nsIs483, nsIs484* | | $P_{gcy\text{-}8}$*:gcy-8:GFP* | This study. Integration of nsEx3945 (Singhvi et al., 2016) |
| *nsIs645, nsIs647* | X | $P_{srtx\text{-}1B}$*:STRX-1:GFP* | This study. Integration of nsEx4078. |
| *nsIs143* | X | $P_{F16F9.3}$*:DsRed* | (Procko et al., 2011) |
| *nsIs109* | | $P_{F16F9.3}$*:DTA(G53E)* | (Bacaj et al., 2008) |
| *dnaIs1, dnaIs2, dnaIs3, dnaIs4* | | $P_{srtx\text{-}1B}$*:HisCl1:SL2:GFP* | This study. Integration of nsEx5340. |
| *dnaIs5* | | $P_{srtx\text{-}1B}$*:SRTX-1:Dendra2* | This study. Integration of dnaEx38. |

| | | | |
|---|---|---|---|
| *dnaIs6-dnaIs9* | | $P_{F53F4.13}$*:CED-10:SL2:mCherry* | This study. Integration of nsEx5365 |

**C. Unstable extra-chromosomal array transgenes and plasmids generated in this study**

All transgenic arrays were generated with 5ng/µl $P_{elt\text{-}2:mCherry}$, 20ng/µl, $P_{mig\text{-}24}$*:Venus*, or 20ng/µL $P_{unc\text{-}122}$*:RFP*(Miyabayashi et al., 1999) as co-injection markers(Abraham et al., 2007; Armenti et al., 2014; Miyabayashi et al., 1999). Further information is available upon request.

| Extra-chromosomal array (nsEx or dnaEX) number | Plasmid | Genotype |
|---|---|---|
| *nsEx3944,nsEx3945, nsEx3946, nsEx3947* | Recombineered fosmid | Singhvi et al, 2016 |
| *nsEx4733, nsEx4734, nsEx4857, nsEx4763* | Recombineered fosmid | $P_{gcy\text{-}18}$*:gcy-18:GFP + elt-2:mCherry* |
| *nsEx4803, nsEx4765* | Recombineered fosmid | $P_{gcy\text{-}23}$*:gcy-23:GFP + elt-2:mCherry* |
| *nsEx4392, nsEx4393, nsEx4394, nsEx4446* | pAS428 | $P_{srtx\text{-}1B}$*:DYF-11:GFP + elt-2:mCherry* |
| *nsEx4051, nsEx4077, nsEx4078* | pAS322 | $P_{srtx\text{-}1B}$*:SRTX-1:GFP* + $P_{unc\text{-}122}$*:RFP* |
| *nsEx4570, nsEx4616, nsEx4688* | pAS447 | $P_{srtx\text{-}1}$*:EGL-1* + $P_{mig\text{-}24}$*:Venus* |

| | | |
|---|---|---|
| *nsEx5266, nsEx5340, nsEx5356* | pAS540 | $P_{srtx-1}$:*HisCl1:SL2:GFP*<br>+ *elt-2:mCherry* |
| *nsEx5365, nsEx5381, nsEx5382* | pAS275 | $P_{F53F4.13}$:*CED-10B:SL2:mCherry*<br>+ $P_{mig-24}$:*Venus* |
| *dnaEx1, dnaEx2, dnaEx3* | pSAR1 | $P_{F53F4.13}$:*CED-12B:SL2:mCherry*<br>+ $P_{unc-122}$:*RFP* |
| *dnaEx19, dnaEx30, dnaEx33* | pSAR7 | $P_{F53F4.13}$:*PSR-1C:SL2:mCherry*<br>+ $P_{unc-122}$:*RFP* |
| *dnaEx29* | pSAR8 | $P_{F53F4.13}$:*CED-10B*$^{G12V}$:*SL2:mCherry*<br>+ $P_{unc-122}$:*RFP* |
| *dnaEx51, dnaEx57, dnaEx59* | pSAR11 | $P_{F53F4.13}$:*CED-10B*$^{T17N}$:*SL2:mCherry*<br>+ $P_{unc-122}$:*RFP* |
| *dnaEx38, dnaEx39, dnaEx40,*<br>*dnaEx41* | pSAR12 | $P_{srtx-1b}$:*SRTX-1:Dendra2*<br>+ $P_{unc-122}$:*RFP* |
| *nsEx5268, nsEx5363, nsEx5380* | pAS247 | $P_{F53F4.13}$:*WSP-1:SL2:mCherry*<br>+ $P_{mig-24}$:*Venus* |

**Plasmids**

CED-10 PLASMIDS: *ced-10*B isoform cDNA was isolated from a mixed stage cDNA library by PCR amplification with primers containing Xma1 and Nhe 1 restriction enzyme sites and directionally ligated into pAS465 ($P_{F53F4.13}$:*SL2:mCherry*)to generate pAS275 plasmid. CED-10$^{G12V}$ and CED-10$^{T17N}$ mutations were derived by site directed mutagenesis of pAS275 plasmid to produce pSAR8 and pSAR11 respectively.

<u>CED-12 PLASMIDS</u>: *ced-12*B isoform cDNA was isolated from a mixed stage cDNA library by PCR amplification with primers containing a XmaI and NheI restriction enzyme sites and directionally ligated into pAS465 to generate the pSAR1 plasmid.

<u>PSR-1 PLASMID</u>: *psr-1* C isoform cDNA was isolated from a mixed stage cDNA library by PCR amplification with primers containing BamHI and NheI restriction enzyme sites, and directionally ligated into pAS465 to generate the pSAR7 plasmid. K324E and K331E mutations were introduced by site directed mutagenesis of pSAR7 to produce pSAR15.

<u>GFP:PSR-1 PLASMID</u>: *psr-1*C isoform cDNA was isolated from a mixed stage cDNA library by PCR amplification with primers containing BamHI and PstI restriction enzyme sites and ligated into pAS516 ($P_{F53F4.13}$*:GFP*) to produce pSAR18.

<u>HisCl1 PLASMID</u>: Histamine gated chloride channel sequence from pNP424(Pokala et al., 2014) was restriction digested with NheI and KpnI enzymes and ligated to pAS178 ($P_{SRTX-1}$*:SL2:GFP*) to produce pAS540.

<u>RECOMBINEERED FOSMIDS</u>: The following fosmids with GFP recombineered in-frame in the coding sequence were obtained from the MPI-TransgeneOme Project: *gcy-8* (Clone ID: 02097061181003035 C08), *gcy-18* (Clone ID: 9735267524753001 E03), *gcy-23* (Clone ID:

6523378417130642 E08).

**Microscopy, Image Processing and Analyses**

Animals were immobilized using either 2mM Tetramizole or 100nm polysterene beads (Bangs Laboratories, Catalog # PS02004). Images were collected on a Deltavision Elite RoHS wide-field deconvolution system with Ultimate Focus(GE), a PlanApo 60x/1.42 NA or OLY 100x/1.40 NA oil-immersion objective and a DV Elite CMOS Camera. Super-resolution microscopy images were collected on the Leica VT-iSIM microscope or the Leica SP8 confocal with Lightning. Images were processed on ImageJ, Adobe Photoshop CC or Adobe Illustrator CC.

Binning categories for population analyses were based on preliminary analyses of population distribution of puncta numbers/animal in wild-type, and mutants with excess puncta (*tax-2*) or reduced puncta mutants (*ced-10, psr-1*). Preliminary analyses of these strains suggested that the bin intervals (0, 1-9 or 10+ puncta) are the most robust, conservative and rapid assessment of phenotypes. Higher than 10 puncta/cell were not readily resolved without post-processing and therefore binned together in population scores. Some genotypes were selected for further *post-hoc* single cell puncta quantification analyses. For this, glia puncta numbers of were quantified using Analyze Particles function in ImageJ on deconvolved images. Individual puncta size measurements were done on yz orthogonal rendering of optical sections using3D objects counter plug-in in ImageJ.

**Electron Microscopy**

Adult hermaphrodites were fixed in 0.8% glutaraldehyde -0.8% osmium tetroxide-0.1 M cacodylate buffer (pH 7.4) for 1 hr at 4°C in the dark and then rinsed quickly several times with 0.1M cacodylate buffer. Animal heads were decapitated and fixed in 1% osmium tetroxide-0.1 M cacodylate buffer overnight at 4°C, quickly rinsed several times in 0.1M Cacaodylate buffer and dehydrated through a graded ethanol series. The samples were then embedded in Eponate 12 resin (Ted Pella, Inc, Redding CA) and polymerized overnight in a 60C oven. 70nm ultrathin serial sections were collected onto pioloform coated slot grids from the anterior tip of the animal to a distance of approximately 7um. Sections were examined on a JEOL 1400 TEM (JEOL, Tokyo, Japan) at an accelerating voltage of 120kV. Images were acquired with a Gatan Rio 4kx4k detector (Gatan, Inc, Pleasanton, CA). Microvilli size measurements were done with ImageJ Measure Function on electron micrograph thin sections.

**Statistical Analyses**

Population puncta scoring was statistically analyzed using Chi Square statistical test in GraphPad Prism 8. Puncta images were quantified using Analyze Particles function in Image J and analyzed with Kruskal-Wallis Test with multiple comparison test in GraphPad Prism 8.

**Chemo-genetic silencing and RNAi**

For chemo-genetic silencing assays, 10mM Histamine (Sigma, Catalog # H7250) was added to NGM agar plates. L4 larval stage transgenic worms expressing HisCl1 in AFD were grown for 24 hours on either normal or Histamine plates and assayed as Day 1 adults (Pokala et al., 2014). Plasmids expressing double-stranded RNA (dsRNA) were obtained from the Ahringer library(Fraser et al., 2000; Kamath, 2003). The L4440 empty vector was used as negative control. RNAi was performed by feeding synchronized L1 animals RNAi bacteria(Timmons, 2004). L4 larva were moved to a fresh plate with RNAi bacteria and scored 24 hours later for glial puncta (*nsIs483)* or AFD-NRE defects (*nsIs645)*.

**Animal Behavior Assays**

Thermotaxis assays were performed on a 18°-26°C linear temperature gradient(Hedgecock and Russell, 1975; Mori and Ohshima, 1995). Animals were synchronized and the staged progeny were tested on the first day of adulthood. Briefly, animals were washed twice with S-Basal and spotted onto the center of a 10-cm plate warmed to room temperature and containing 12 mL of NGM agar. The plate was placed onto the temperature gradient (17-26$^{o}$C) with the addition of 5 mL glycerol to its bottom to improve thermal conductivity. At the end of 45 mins, the plate was inverted over chloroform to kill the animals and allowing easy counting of animals in each bin. The plates have an imprinted 6x6 square pattern which formed the basis of the 6 temperature bins. Each data point is the average of 3-8 assays with ~150 worms/assay.

**Fig. S1.**

**AMsh glia engulf AFD-NRE fragments. (A)** AFD-NRE labeled fragments observed across transgenic animal strains carrying different promoters or protein tags. X=axis: genotype; Y-axis: percent animals with AFD-NRE labeled puncta inside AMsh soma. N= number of animals analyzed. **(B)** Time-stamped stills from movie (Supplemental S1) of AFD-NRE dissociation of fragments. Each colored arrowhead tracks an individual fragment moving away from AFD-NRE. Scale bar: 5µm. **(C)** Quantification of average puncta diameter within AMsh glial cell soma **(D)** Quantification average AFD-NRE microvilli diameter from electron micrographs **(E)** Electron micrograph through AFD-NRE microvilli of an animal. An individual microvillum taken for diameter measurement in Fig 1E is outlined in yellow. Scale bar: 500nm **(F,G)** Quantification of puncta in ipsi- and contra-lateral AMsh glial cell soma with AFD neurons ablated by laser (F) or genetically (G). N= number of animals assayed.

#### SUPPLEMENTARY FIGURE 1

Raiders et al

A

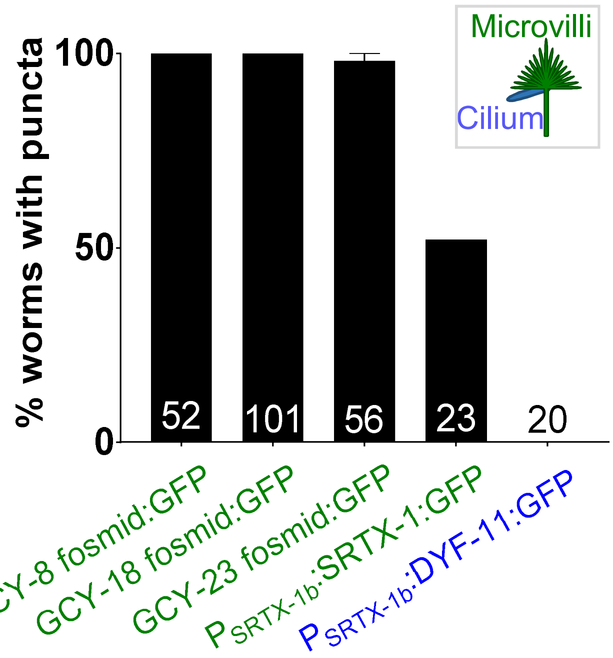


B

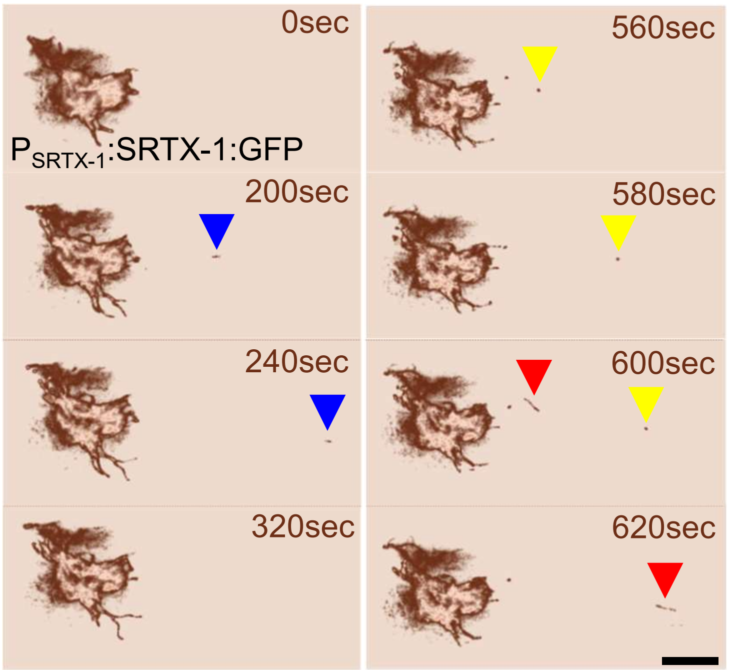


C

| Puncta average diameter (nm) | Average Deviation | N (animals) |
|---|---|---|
| 541 | 145 | 7 |

D

| Microvilli average diameter (nm) | Average Deviation | N (animals) |
|---|---|---|
| 214 | 30 | 3 |

E

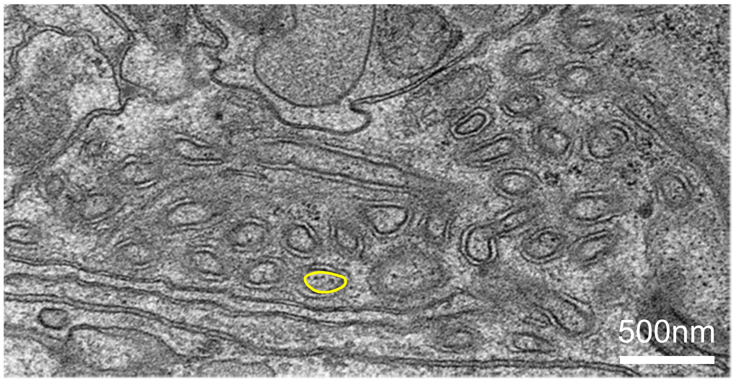


F

| **Laser ablation** in *nsIs483* | % animals with ipsilateral puncta | % animals with contralateral puncta | Total animals |
|---|---|---|---|
| Mock ablation | 100 | 100 | 58 |
| AFD ablation | 0 | 100 | 24 |

G

| **Genetic ablation** in *nsIs483; nsEx4616[$P_{SRTX-1}$:EGL-1]* | % animals with ipsilateral puncta | % animals with contralateral puncta | Total animals |
|---|---|---|---|
| AFD present | 100 | 100 | 130 |
| AFD absent | 0 | 100 | 21 |

**Fig. S2.**

**AMsh glia engulf AFD-NRE microvilli in activity-dependent manner.**

**(A)** Fluorescence and DIC micrographs showing expression of ciliary DYF-11:GFP under an AFD neuron-specific promoter in AFD cilia. C (arrowhead), cilia. D (arrow), AFD dendrite. **(B)** Average number of fragments in animals cultivated at 15, 20, or 25°C. Refer Figure 1I for data presentation details. Median puncta counts and N (number of animals): 15°C (6±2 puncta, n=8 animals) , 20°C (15±1 puncta, n=54 animals), 25°C (27±3 puncta, n=16 animals) **(C)** Population counts of animals with AMsh glial puncta in animals with RNAi (control, *pat-2)* in *tax-2(p691)* (A) or *tax-2(p691); psr-1(tm469)* mutant (B) animals. Refer Figure 1H for data presentation details.

### SUPPLEMENTARY FIGURE 2



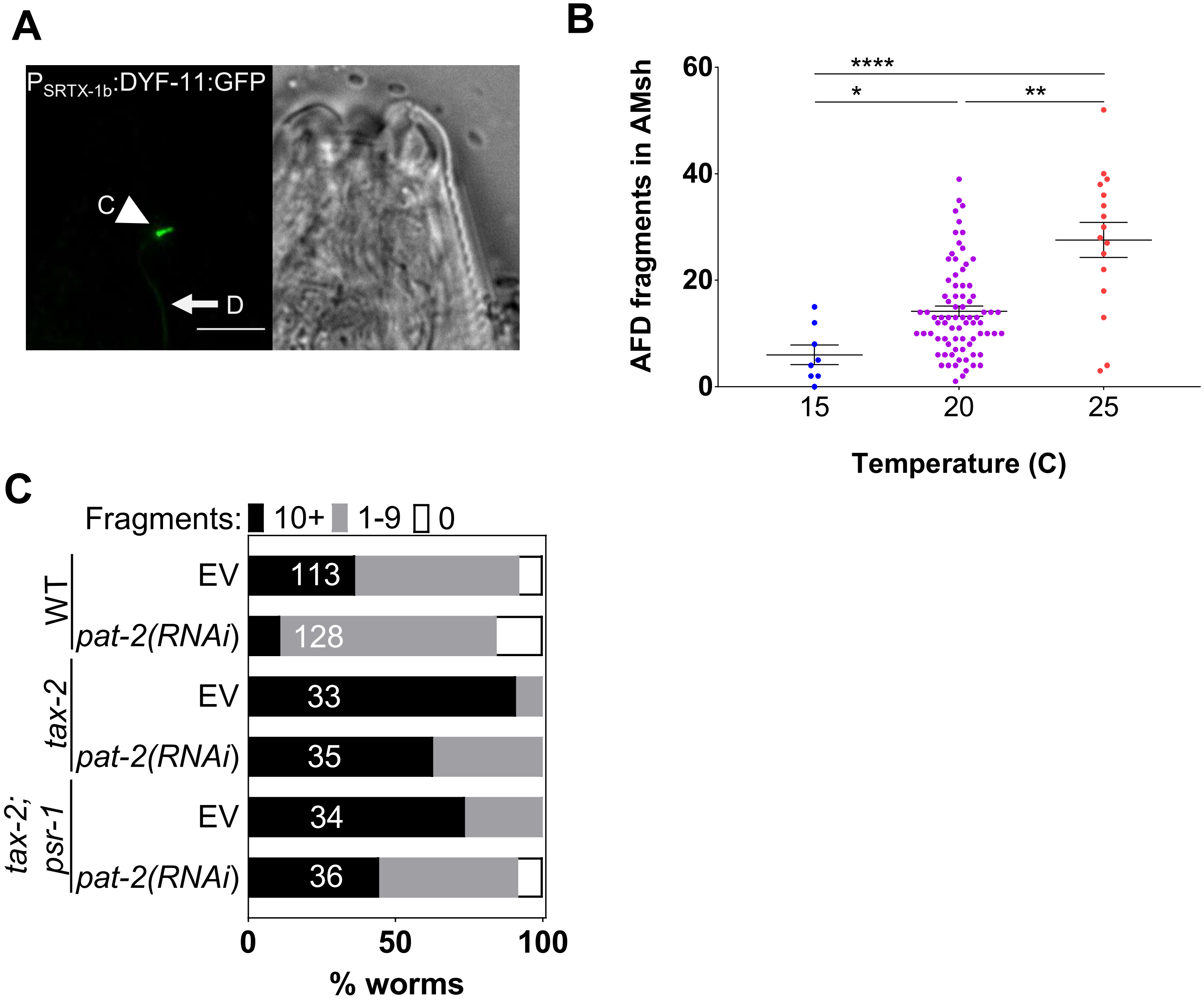

**Fig. S3.**

**AMsh glia engulf AFD-NRE by repurposing components of the phagocytosis machinery. (A)** Schematic of the genetic pathway underlying apoptotic corpse engulfment in *C. elegans*. **(B)** Percent animals with AMsh glial puncta in genetic backgrounds indicated. N= number of animals scored. Refer Extended Figure 1B for data presentation details.

**SUPPLEMENTARY FIGURE 3** Raiders et al

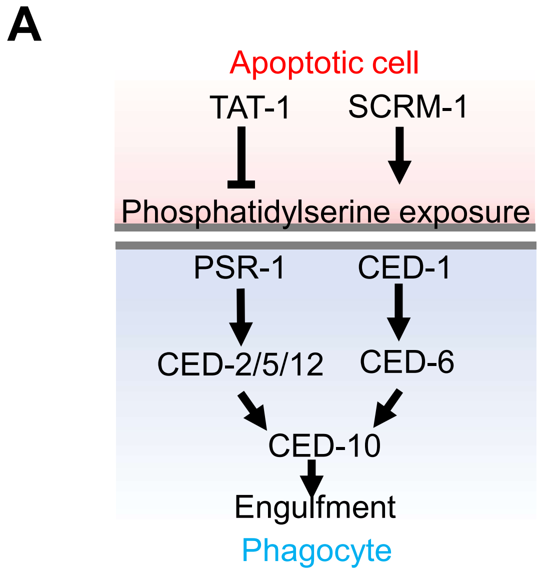


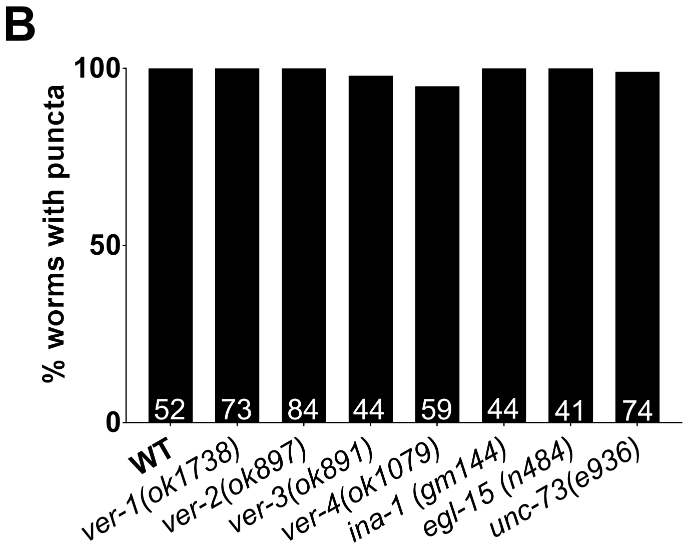

**Fig. S4.**

**Activation of CED-10/Rac1 regulates AFD NRE shape (A)** Day 1 AFD NRE defects in animals expressing constitutive active CED-$10^{G12V}$ in AMsh glia. **(B)** Proportion of worms with defective AFD-NRE shape on Day 1 and 3 of adulthood in animals expressing constitutive active CED-$10^{G12V}$ or dominant negative CED-$10^{T17N}$.

### SUPPLEMENTARY FIGURE 4

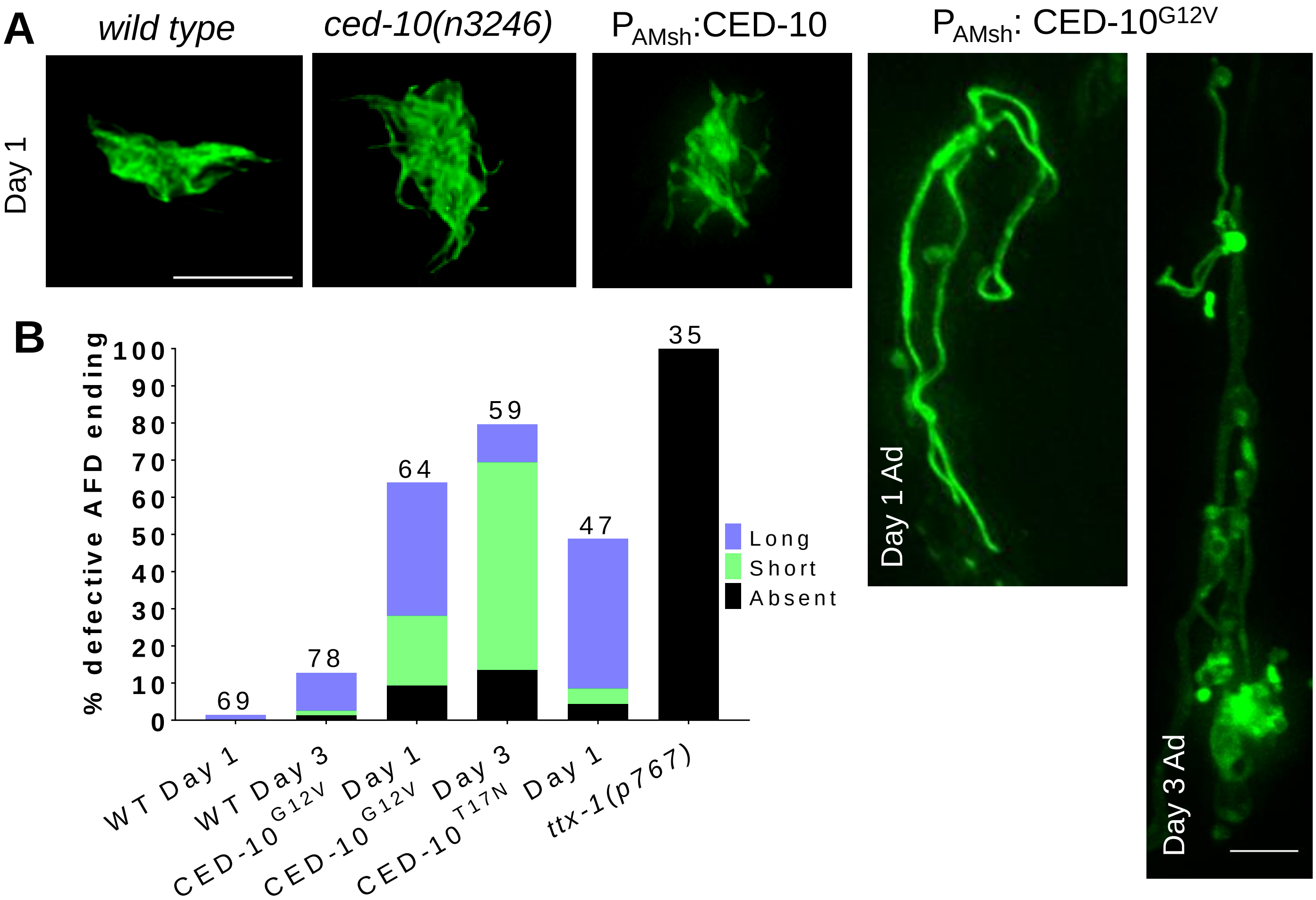

**Movie S1.**

**Dissociation of AFD-NRE fragments.**

Movie of an animal's AFD-NRE, labeled with GFP and imaged *in vivo* at 7 frames/second, shows fragments blebbing at regular intervals.

**SUPPLEMENTARY MATERIALS REFERENCES**


Abraham, M.C., Lu, Y., and Shaham, S. (2007). A morphologically conserved nonapoptotic program promotes linker cell death in Caenorhabditis elegans. Dev Cell *12*, 73-86.

Armenti, S.T., Lohmer, L.L., Sherwood, D.R., and Nance, J. (2014). Repurposing an endogenous degradation system for rapid and targeted depletion of C. elegans proteins. Development *141*, 4640-4647.

Bacaj, T., Tevlin, M., Lu, Y., and Shaham, S. (2008). Glia are essential for sensory organ function in C. elegans. Science *322*, 744-747.

Colosimo, M.E., Brown, A., Mukhopadhyay, S., Gabel, C., Lanjuin, A.E., Samuel, A.D., and Sengupta, P. (2004). Identification of thermosensory and olfactory neuron-specific genes via expression profiling of single neuron types. Curr Biol *14*, 2245-2251.

Fraser, A.G., Kamath, R.S., Zipperlen, P., Martinez-Campos, M., Sohrmann, M., and Ahringer, J. (2000). Functional genomic analysis of C. elegans chromosome I by systematic RNA interference. Nature *408*, 325-330.

Hedgecock, E.M., and Russell, R.L. (1975). Normal and mutant thermotaxis in the nematode Caenorhabditis elegans. Proc Natl Acad Sci U S A *72*, 4061-4065.

Kamath, R. (2003). Genome-wide RNAi screening in Caenorhabditis elegans. Methods *30*, 313-321.

Miyabayashi, T., Palfreyman, M.T., Sluder, A.E., Slack, F., and Sengupta, P. (1999). Expression and function of members of a divergent nuclear receptor family in Caenorhabditis elegans. Dev Biol *215*, 314-331.

Mori, I., and Ohshima, Y. (1995). Neural regulation of thermotaxis in Caenorhabditis elegans. Nature *376*, 344-348.

Pokala, N., Liu, Q., Gordus, A., and Bargmann, C.I. (2014). Inducible and titratable silencing of Caenorhabditis elegans neurons in vivo with histamine-gated chloride channels. Proc Natl Acad Sci U S A *111*, 2770-2775.

Procko, C., Lu, Y., and Shaham, S. (2011). Glia delimit shape changes of sensory neuron receptive endings in C. elegans. Development *138*, 1371-1381.

Singhvi, A., Liu, B., Friedman, C.J., Fong, J., Lu, Y., Huang, X.Y., and Shaham, S. (2016). A Glial K/Cl Transporter Controls Neuronal Receptive Ending Shape by Chloride Inhibition of an rGC. Cell *165*, 936-948.

Timmons, L. (2004). Endogenous inhibitors of RNA interference in Caenorhabditis elegans. Bioessays *26*, 715-718.
